# Single-molecule architecture and heterogeneity of human telomeric DNA and chromatin

**DOI:** 10.1101/2022.05.09.491186

**Authors:** Danilo Dubocanin, Adriana E. Sedeno Cortes, Jane Ranchalis, Taylor Real, Ben Mallory, Andrew B. Stergachis

## Abstract

Telomeres are essential for linear genomes, yet their repetitive DNA content and somatic variability has hindered attempts to delineate their chromatin architectures. We performed single-molecule chromatin fiber sequencing (Fiber-seq) on human cells with a fully resolved genome, enabling nucleotide-precise maps of the genetic and chromatin structure of all telomeres. Telomere fibers are predominantly comprised of three distinct chromatin domains that co-occupy individual DNA molecules – multi-kilobase telomeric caps, highly accessible telomeric-subtelomeric boundary elements, and subtelomeric heterochromatin. Extended G-rich telomere variant repeats (TVRs) punctuate nearly all telomeres, and telomere caps imprecisely bridge these degenerate repeats. Telomeres demonstrate pervasive somatic alterations in length, sequence, and chromatin composition, with TVRs and adjacent CTCF-bound promoters impacting their stability and composition. Our results detail the structure and function of human telomeres.

**One sentence summary:** We use single-molecule chromatin fiber sequencing to detail the structure and function of human telomeric DNA and chromatin.

## Main

Human telomeres are composed of tandem TTAGGG DNA repeat arrays^1^, which are encased by a telomere cap chromatin structure via the recruitment of specialized sequence-specific shelterin protein complex members^2^. Despite the protective role of telomere caps in maintaining the stability of telomere repeat arrays, telomeres are prone to undergo both germline and somatic genetic alterations^3, 4^, with telomere shortening playing a central role in various human diseases and normal aging processes^5^. The highly repetitive DNA content and somatic variability within telomeres has curtailed the use of sequencing-based approaches for interrogating their genetic and chromatin structure. Specifically, accurate reference sequences for telomeres are largely unavailable, and short-read based chromatin profiling methods such as DNaseI-seq^6, 7^, ATAC-seq^8^, or ChIP-seq^9^ cannot disentangle single-molecule chromatin architectures across highly-repetitive and somatically-variable multi-kilobase sequences.

Consequently, our understanding of the basic structure and function of the DNA and chromatin component of telomeres is quite limited. For example, proximal telomeric sequences exposed via telomere shortening have been shown to frequently harbor telomere variant repeats (TVRs) comprised of degenerate non-TTAGGG DNA repeats^4, 10^, yet the impact of these degenerate repeats on telomere cap formation is largely unknown. In addition, subtelomeric regions are populated by diverse chromatin features, such as CpG islands that promote TElomeric Repeat-containing RNA (TERRA) transcription^11, 12^, CCCTC-binding factor (CTCF)-occupied regulatory elements^13, 14^, and constitutive heterochromatin domains^15, 16^. However, the interplay of these chromatin domains along individual DNA molecules is unresolved, and their potential impact on telomere structure and function is largely unknown.

The completion of the first complete human reference genome from CHM13 cells^17^ opens the possibility of studying the DNA and chromatin architecture across all human telomeres. Furthermore, emerging long-read sequencing-based chromatin profiling methods have the potential to overcome the inherent mapping limitations of short-read based chromatin methods at repetitive DNA elements^18–20^ (**fig. S1A**). For example, single-molecule chromatin fiber sequencing (Fiber-seq) is capable of mapping nucleotide-precise patterns of chromatin accessibility, transcription factor (TF) occupancy, and nucleosome positioning along multi-kilobase chromatin fibers^18^, and we have optimized the original protocol for mapping architectures of human chromatin (**fig. S2**). Specifically, Fiber-seq utilizes a non-specific DNA N^6^-adenine methyltransferase (m6A-MTase)^21^ to stencil the chromatin architecture of individual multi-kilobase fibers onto their underlying DNA templates via methylated adenines (**Fig. 1A**), which is a non-endogenous DNA modification in humans^22, 23^. The genetic and chromatin architecture of each fiber is directly read using highly accurate single-molecule Pacific Biosciences (PacBio) HiFi long-read DNA sequencing, which is capable of accurately distinguishing m6A-modified bases^18^. Given the high sequence accuracy and long read lengths obtained during Fiber-seq, as well as the evenly distributed A/T content of telomeric DNA (**fig. S1B**), we hypothesized that Fiber-seq could uniquely enable the interrogation of the genetic and chromatin architectures within human telomeres at single-molecule and single-base resolution.

**Fig. 1.**
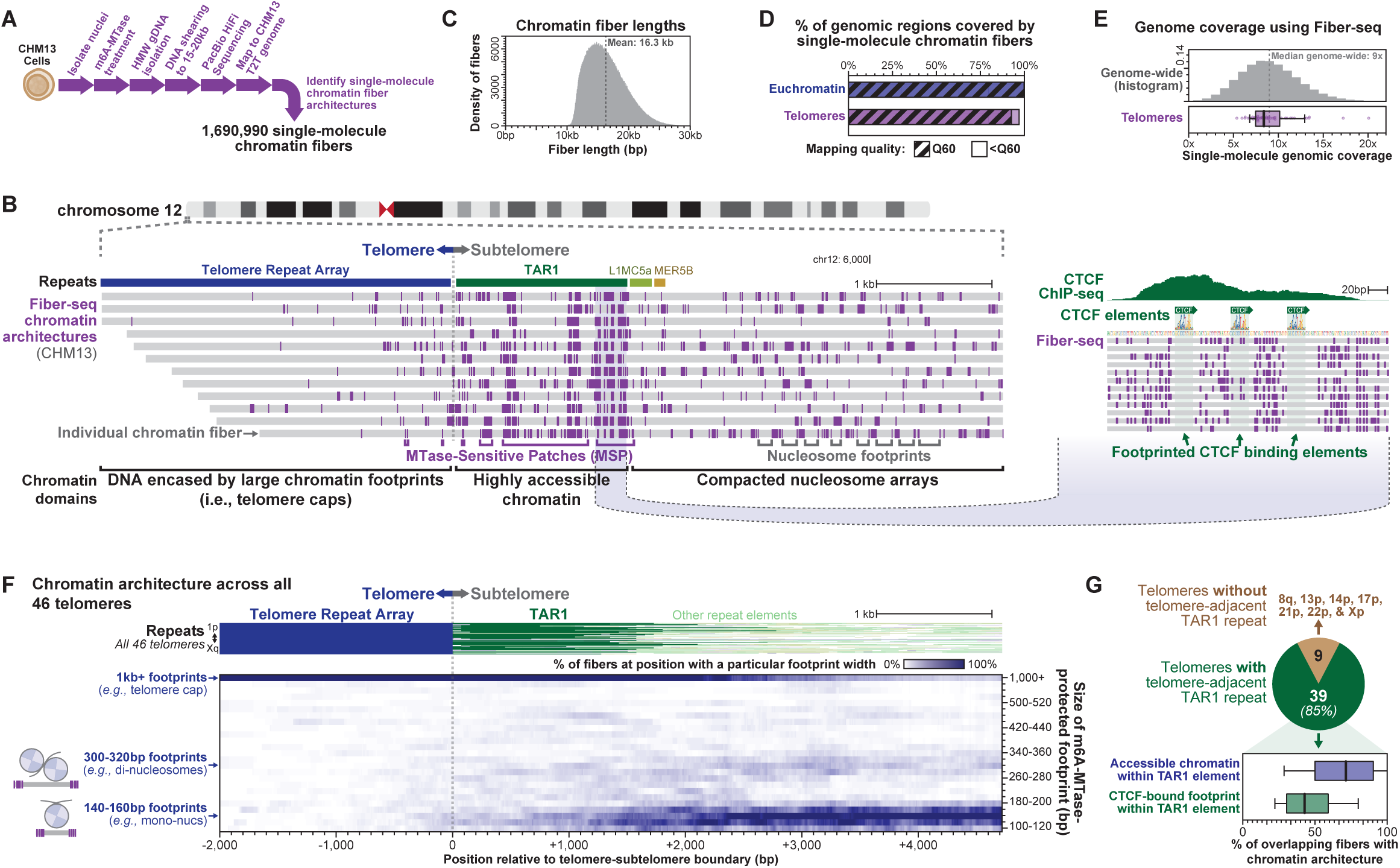
Single-molecule chromatin fiber sequencing of all human telomeres. **(A)** Fiber-seq schematic for mapping chromatin architectures from a complete human telomere-to- telomere epigenome. **(B)** Genomic locus showing per-molecule Fiber-seq chromatin architectures for all fibers overlapping the chromosome 12p telomere. Location of repeat elements indicated above. (right) Insert of TAR1 repeat, highlighting CTCF binding elements and single-molecule CTCF occupancy events. **(C)** Histogram showing the size distribution of chromatin fibers sequenced using Fiber-seq. **(D)** Mapping quality of the 1.7 million chromatin fibers delineated using Fiber-seq as well as the percentage of all euchromatic and telomeric bases in the CHM13 genome covered by Fiber-seq data. **(E)** Coverage of single-molecule chromatin fibers (top) genome-wide or (bottom) along individual telomeres (including 25kb subtelomeric region). **(F)** Heatmap showing the distribution of footprint sizes relative to the telomere-subtelomere boundary across all CHM13 telomeres. (Top) Repeat elements present across all 46 chromosome ends. **(G)** Proportion of telomeres that contain telomere-adjacent TAR1 repeats, as well as the single-molecule chromatin accessibility and CTCF occupancy at these repeats.

### Mapping chromatin architectures of human telomeres

To investigate the chromatin architecture of human telomeres, we applied Fiber-seq to the human hydatidiform mole CHM13_hTERT_ cell line (hereafter referred to as CHM13)^24^, the same line used for constructing the first human telomere-to-telomere (T2T) reference genome^17, 25^, as well as the human lymphoblastoid cell line GM12878 as a comparison. By coupling a fully assembled genome map with matched long-read epigenome data, we were able to overcome the substantial genetic heterogeneity of these regions^26^ that appears to be enriched within these highly repetitive genomic regions (**fig. S3A**).

Application of Fiber-seq to CHM13 cells resulted in 1.7 million individual ∼16kb chromatin fiber architectures (**Figs. 1, A to D and figs. S3, B to D**), enabling the nucleotide-precise delineation of over 230 million chromatin features, such as nucleosomes, accessible regulatory elements, and single-molecule TF binding events (**Figs. 1, B and C**). Furthermore, the high sequence accuracy of these reads enabled us to uniquely map the chromatin architecture of over 97% of the human genome, including all telomeres (**Figs. 1, D and E**).

### Human telomeres contain three distinct chromatin domains

We first investigated the single-molecule chromatin organization overlying telomeres and subtelomeres, which are known to contain diverse and somewhat contradictory chromatin features^2, 11–16^. Single-molecule chromatin fiber sequencing of human telomeres exposed that telomeric and subtelomeric chromatin is predominantly segmented into three distinct chromatin domains: (1) an MTase-protected telomere cap structure occupying the telomere repeat array; (2) a highly accessible subtelomeric chromatin domain immediately adjacent to most telomere repeat arrays; and (3) an extended heterochromatic subtelomeric region (**Figs. 1, B and F; fig. S4; fig. S5**). Notably, the highly accessible subtelomeric chromatin domain localizes to telomere-adjacent Telomere-Associated Repeat (TAR1) repeats^27^, which bookend 85% of all telomeres in CHM13 cells and are frequently occupied by CTCF elements (**Figs. 1F and G**). Overall, 52% of telomeric fibers contain all three of these domains abutting each other within ∼5 kb – indicating that previously observed telomeric and subtelomeric chromatin features are physically partitioned across these three distinct chromatin domains.

### Telomere caps heterogeneously encase telomere repeat arrays

Although nucleosomes appear capable of occupying telomere repeat arrays (**Figs. 1F and fig. S6A**), telomere repeat arrays are largely occluded from m6A-MTase methylation, producing extended chromatin footprints along the terminal portion of each telomere (i.e., telomere caps) (**Fig. 1F; Figs. 2, A and B**). The transition between these extended telomere cap chromatin footprints and their neighboring nucleosome-demarcated chromatin domain is highly dynamic, varying by over 1 kb between individual fibers from the same telomere (**Fig. 2C**), indicating that telomere cap formation is highly heterogeneous on a per-molecule level.

**Fig. 2.**
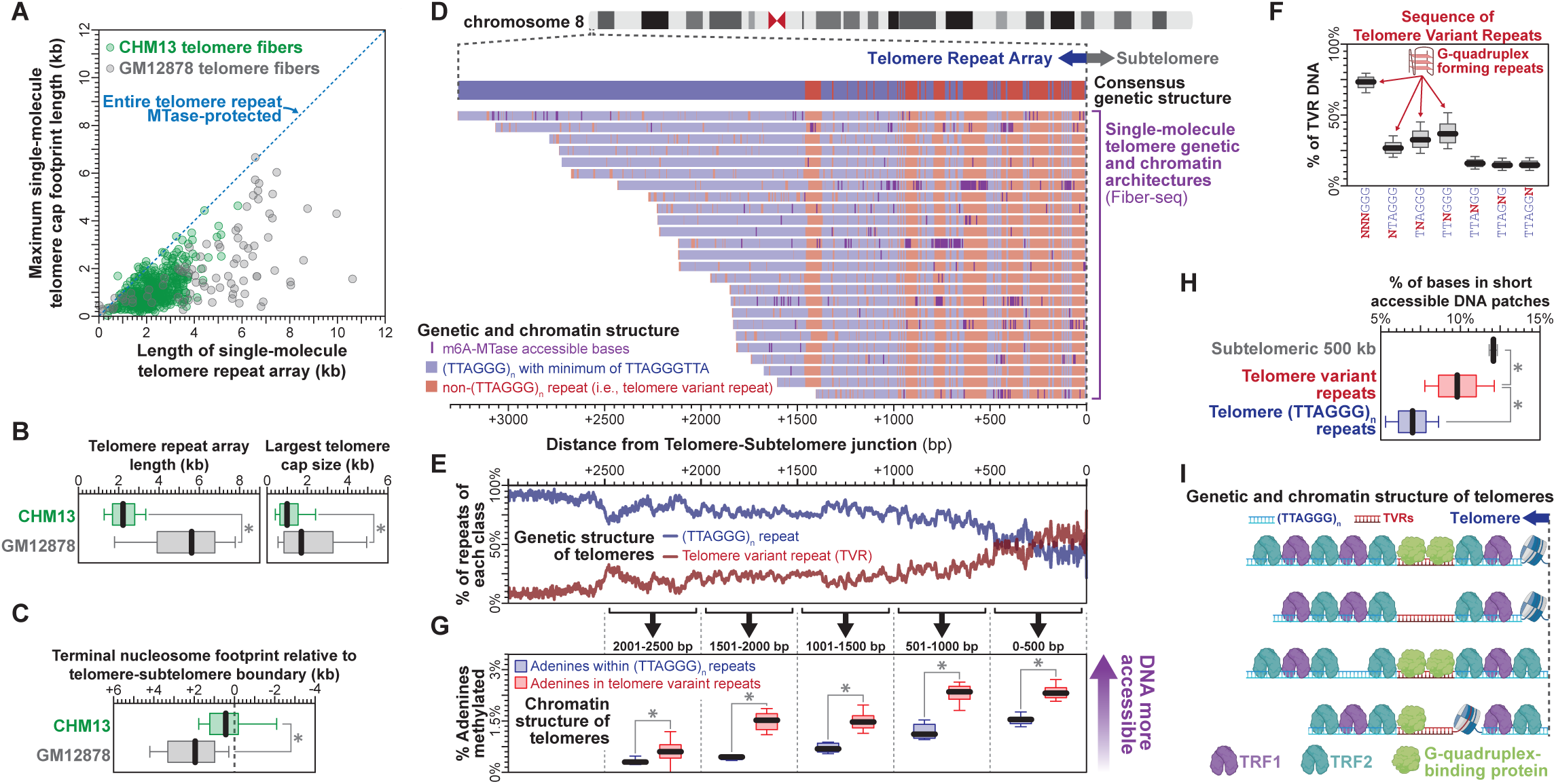
Telomeric cap architecture of human telomeres. **(A)** The association between per-molecule telomere length and max telomere cap length across all telomeres from CHM13 and GM12878 cells. **(B)** Box-and-whisker plots showing (left) per-fiber telomere lengths, and (right) per-fiber telomere largest cap sizes between telomere fibers from CHM13 and GM12878 cells (* p-value <0.01 Mann-Whitney). **(C)** Box-and-whisker plots showing location of terminal nucleosome footprints between telomere fibers from CHM13 and GM12878 cells (* p-value <0.01 Mann-Whitney). **(D)** Genomic locus showing per-molecule genetic and chromatin architectures for all fibers overlapping the chromosome 8p telomere. Location of per-fiber TTAGGG, and non-TTAGGG TVRs in blue and red respectively. Consensus sequence of telomere shown above. **(E)** Plot showing the percentage of telomeres containing TTAGGG repeats or TVRs at each base relative to telomere- subtelomere boundary. **(F)** Sequence of TVRs within telomeric DNA. **(G)** m6A-methylation frequency of TTAGGG repeats and TVRs within telomeres, relative to telomere-subtelomere boundary. **(H)** Localization of short accessible patches of DNA within TTAGGG repeats and TVRs, as well as subtelomeric regions (* p-value <0.01 Mann-Whitney). **(I)** Model for the formation of telomeric cap structures across telomere repeat arrays.

Notably, we observed that interstitial TTAGGG repeats similarly form extended chromatin footprint structures, albeit to a slightly less degree (**fig. S6B**). As interstitial TTAGGG repeats are known to recruit shelterin proteins^28–30^, this finding further supports the role of shelterin in forming these extended chromatin footprints along telomere repeat arrays.

### Short telomere repeat arrays form short telomere caps

To determine the impact of the length of the telomere repeat array on telomere cap size, we compared telomeric chromatin architectures across CHM13 cells and GM12878 cells, as GM12878 cells have markedly longer telomere repeat arrays (**Figs. 2, A and B**). Notably, shorter telomere repeat arrays were associated with shorter overlying telomere caps, with telomere caps almost always terminating at or before the telomere-subtelomere boundary (**Figs. 2, A and B**). In addition, telomere caps along multi-kilobase telomere repeat arrays were often fragmented into multiple parts by short MTase-accessible patches and nucleosome footprints (**Fig. 2A; figs. S7, A and B**), further highlighting the impact of telomere repeat array length on overlying chromatin cap formation.

### Telomere caps imprecisely bridge telomere variant repeats

Telomere caps are known to be formed by the shelterin complex^31^, which interacts with telomere repeat sequences through the sequence-specific binding of TRF1 and TRF2 to TTAGGGTT sequences^28^.

However, although human telomeres are predominantly composed of TTAGGG repeats, non-TTAGGG telomere variant repeats (TVRs) are known to localize to the proximal end of human telomeres^4, 10^, raising questions about the potential impact of these TVRs on telomere cap formation and function. We observed that the proximal end of telomere repeat arrays are pervasively punctuated by TVRs across nearly all human telomeres in CHM13 cells (**Figs. 2, D and E; fig. S7B**). These TVRs frequently extend to several hundred bases in length and are often stably present across individual fibers from the same telomere. Notably, the sequence of these TVRs largely maintains the GGG triplet (**Fig. 2F**), indicating that although these repeats lack TRF1/TRF2 binding elements, they still maintain the potential for forming G- quadraplex DNA – a core feature of TTAGGG repeats.

We find that TVRs show significantly higher MTase-accessibility when compared to their neighboring canonical TTAGGG repeats (**Fig. 2G and fig. S7C**) and are significantly more likely to harbor short MTase-accessible patches of chromatin (**Fig. 2H**). However, most telomere cap structures extend through TVRs (**Fig. 2D and fig. S7B**), indicating that the majority of TVRs remain encased by proteins *in vivo*. Overall, these findings demonstrate that although telomere caps are prone to be disrupted by TVRs, telomere caps nonetheless often bridge TVRs despite these degenerate repeats lacking TRF1/TRF2-binding elements (**Fig. 2I**).

### Somatic stability of telomere variant repeats

Given the pervasive punctuation of telomere repeat arrays with TVRs, as well as their potential impact on telomere cap function, we next sought to determine whether TVRs are associated with altered telomere somatic stability. Notably, although individual fibers arising from the same telomere often vary by several kilobases in length^32^, only 5.0% of telomeres were shortened beyond their terminal TVR (**Fig. 3A**), indicating that stable variation in telomere fiber lengths is largely limited to the sequence distal to these terminal TVRs.

**Fig. 3.**
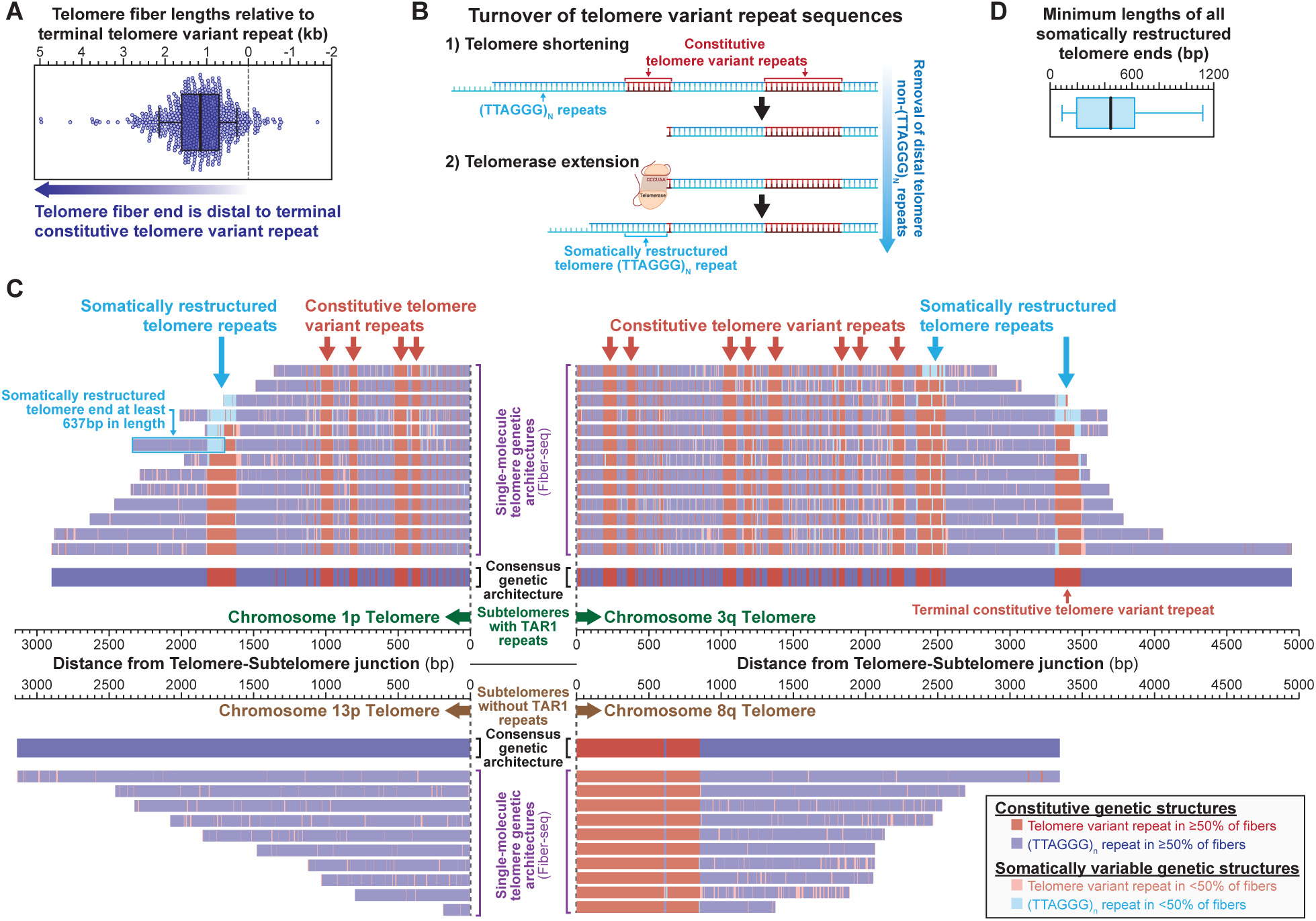
Somatic stability of telomere variant repeats. **(A)** Swarm and box-and-whisker plots showing the length of each telomere fiber from CHM13 cells relative to the most distal constitutive TVR on that telomere. **(B)** Model showing the impact of telomere shortening and telomerase lengthening on the somatic preservation of TVRs. **(C)** Genomic loci showing per-molecule genetic architectures for all fibers overlapping the chromosome 1p, 13p, 3q, and 8q telomeres. Consensus sequences for each telomere are shown in middle. Telomere fibers colored based on their sequence relative to the consensus sequence, with repeats matching the consensus in dark and repeats not matching the consensus in light colors. TTAGGG repeats, and TVRs on each fiber colored in blue and red respectively. **(D)** Box-and-whisker plot showing the minimum lengths of somatically restructured repeats across all 46 CHM13 telomeres.

As CHM13 cells express telomerase, telomeres should be in a steady state balance between shortening and elongation, with newly synthesized telomere DNA predominantly comprised of TTAGGG repeats. As such, if telomere shortening past these terminal TVRs is compatible with cell survival, we would frequently observe telomere fibers with their terminal TVRs converted into canonical TTAGGG repeats (**Fig. 3B**). However, only 3% of telomere fibers exhibited patterns consistent with having their terminal TVR somatically restructured into TTAGGG repeats (**Fig. 3C and fig. S8**). Furthermore, these somatically restructured repeats appear only in the terminal ∼400bp of each telomere, suggesting that telomere fiber shortening beyond this point is not stably maintained in CHM13 cells^33^.

### Stereotyped CTCF-bound TAR1 promoters bookend most telomeres

Next, we sought to characterize the sequence and chromatin content of telomere-adjacent subtelomeric regions. Some of these regions are known to contain CTCF binding elements and CpG islands essential for initiating transcription of the long noncoding telomeric repeat RNA TERRA^12^, which plays a role in maintaining telomere function and stability^13, 34, 35^. We observed that 39 of the 46 telomeres in CHM13 cells harbored telomere-adjacent TAR1 repeats, which were almost uniformly punctuated by a highly accessible chromatin domain marked with CTCF-bound elements and CpG island promoters (**Fig. 1G; Fig. 4A; fig. S9**), consistent with previously identified TERRA promoters^11^. The chromatin structure of telomere-adjacent TAR1 repeats is highly stereotyped, with CTCF occupancy precisely demarcating the boundary between a highly accessible CpG island and adjacent compacted subtelomeric heterochromatin (**Fig. 1B**). Notably, TAR1 repeats almost uniformly contain 2+ CTCF proteins bound simultaneously in tandem along the same molecule of DNA (**Fig. 4, B and C**) with their N-terminal cohesin interacting domains oriented away from the telomere (**Fig. 4, B and D**) – a configuration that should promote insulation of the TAR1 CpG island from subtelomeric heterochromatin^36^ and suppress antisense transcription from the TAR1 CpG island that is directed away from the telomere^37^.

**Fig. 4.**
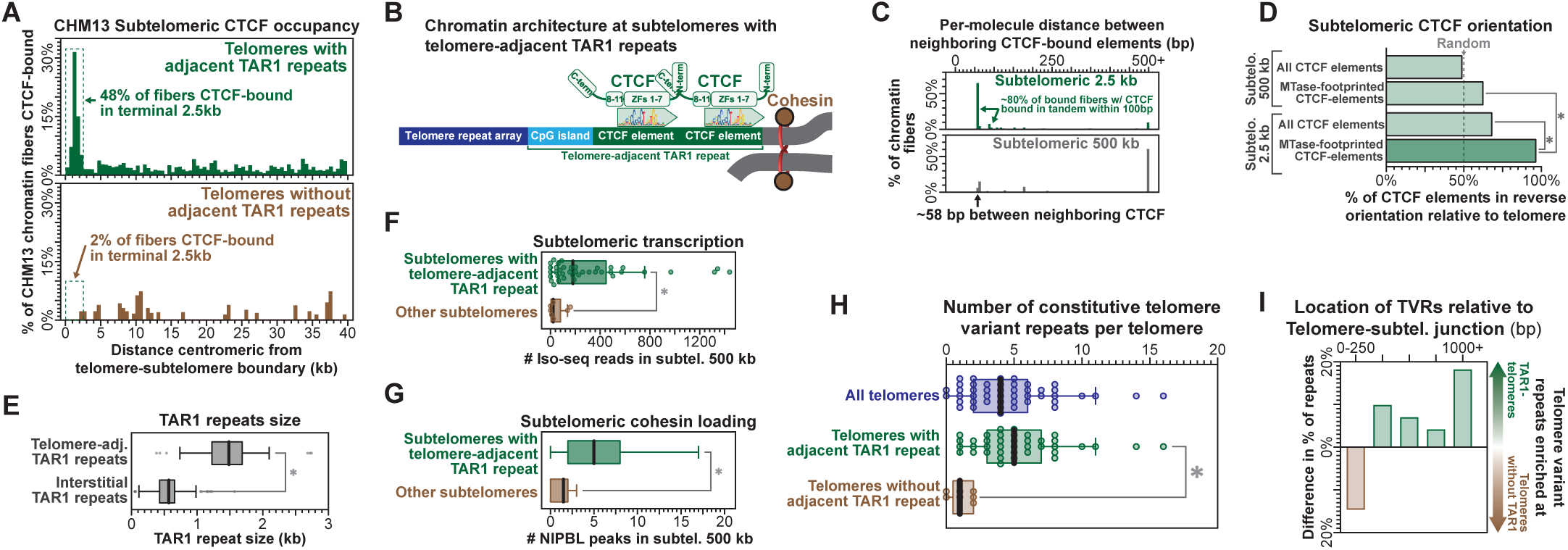
Telomere-adjacent TAR1 repeats impact telomere stability and composition. **(A)** The percentage of fibers that contain a CTCF-footprinted element within 500 bp bins relative to the telomere-subtelomere boundary. Telomeres (top) with and (bottom) without a telomere-adjacent TAR1 repeat. **(B)** Model for CTCF occupancy at telomere-adjacent TAR1 repeats. **(C)** Distance between neighboring MTase-footprinted CTCF elements along the same molecule, by location of footprint relative to telomere-subtelomere boundary. **(D)** The percentage of all CTCF elements, as well as MTase- footprinted CTCF elements, that are oriented in the reverse orientation relative to the telomere (* p- value <0.01 Z test). **(E)** Box-and-whisker plot of TAR1 repeat size for telomere-adjacent and interstitial TAR1 repeats. **(F,G)** Total number of (F) Iso-seq reads or (G) NIPBL ChIP-seq peaks within the 500 kb subtelomeric region for telomeres with or without a telomere-adjacent TAR1 repeat (* p-value <.01 Mann-Whitney). **(H)** Number of constitutive TVRs within telomeres with or without a telomere-adjacent TAR1 repeat (* p-value <.01 Mann-Whitney). **(I)** Location of TVRs within telomeres relative to telomere- subtelomere boundary for telomeres with or without a telomere-adjacent TAR1 repeat.

TAR1 repeats also localized to numerous interstitial locations throughout the genome, but most interstitial TAR1 repeats are fragmented and lack chromatin accessibility and CTCF occupancy (**Fig. 4E; fig. S10, A and B**). However, intact interstitial TAR1 repeats often form gene promoters via their encompassed CpG island (**fig. S10B**). For example, the telomere-to-telomere fusion of the ancestral chromosome 2 at 2q14.1^38^ results in an interstitial TAR1 repeat that has been coopted as a promoter for the gene *DDX11L2* (**fig. S10C**), further demonstrating that TAR1 repeats can form directional gene promoters.

### Telomere-adjacent TAR1 repeats promote TVR stability

Telomeres without adjacent TAR1 repeats preferentially have transcriptionally inactive subtelomeric regions (**Fig. 4F**) that lack cohesin loading (**Fig. 4G and fig. S11**), suggesting that TAR1 repeats may play a unique role in guiding telomere function and stability at transcriptionally active chromosome ends.

Comparing the stability of telomeres with and without TAR1 repeats revealed that overall, telomeres with adjacent TAR1 repeats are significantly longer and are modestly more variable the length (**fig. S12**), indicating that the steady-state balance between telomere shortening and elongation is significantly impacted by TAR1 repeats. Furthermore, telomeres with and without adjacent TAR1 repeats markedly diverged in their sequence composition. Specifically, CHM13 telomeres without adjacent TAR1 repeats contained few, if any, blocks of TVRs (**Fig. 3C and Fig. 4H**), and when present these TVRs were limited to the proximal region of each telomere (**Fig. 4I**). Together, these findings demonstrate that one of the dominant features of telomere-adjacent TAR1 repeats is the stabilization of TVRs within the telomere.

## Discussion

We identify highly stereotyped single-molecule chromatin and genetic architectures across all human telomeres, finding that the majority of human telomere fibers are punctuated by three distinct chromatin domains that physically co-occupy the same fiber. This finding demonstrates that previously observed diverse telomeric and subtelomeric chromatin features^11–16^ are physically partitioned on a per-molecule basis.

We delineate the architecture of telomere caps at single-nucleotide and single-molecule resolution, demonstrating that nucleosome-demarcated chromatin transitions into telomere caps within the proximal ∼500bp of each telomere fiber. However, this transition is highly heterogeneous on a per-molecule level. These single-molecule chromatin measurements contrast prior bulk studies that suggest pervasive nucleosome occupancy throughout the telomere repeat arrays^39, 40^. It is likely that this discrepancy results from bulk averaging of rare nucleosome occupancy events across telomere repeat arrays, as large telomere caps often appear fragmented into multiple sections that can be bookended by nucleosome sized footprints. Alternatively, it is also possible that telomere caps themselves include nucleosomes, with the entry/exit sites of nucleosomes being fully protected by shelterin complex members^41^ such that nucleosome-bound DNA that might otherwise undergo stochastic site-exposure^42^ is sheltered from m6A-MTase activity.

We demonstrate pervasive somatic alterations in the length, sequence, and chromatin composition of telomeres. Specifically, telomeres appear pervasively punctuated by non-TTAGGG TVRs that are subject to somatic restructuring through the action of telomerase during the course of telomere shortening and elongation. Notably, TVRs impact the ability of telomeres to faithfully form telomere caps, with telomere caps imprecisely bridging these degenerate sequences. Importantly, TVRs preferentially maintain the GGG triplet, indicating that they are still capable of forming G-quadraplex DNA. As such, although TVRs lack binding sequences for the shelterin complex members TRF1/TRF2^28^, it is likely that they are being bridged by additional proteins that directly interact with both G-quadraplex DNA and shelterin complex members^43^.

Our findings also demonstrate TAR1 repeats as a functional repeat that establishes a highly structured and accessible directional promoter element at the boundary between telomeric and subtelomeric DNA on 85% of CHM13 telomeres. Notably, these telomere-adjacent TAR1 repeats are associated with the stabilization of TVRs within the telomere. However, the exact mechanism by which they accomplish this remains unknown. Similarly, the functional significance of having 7 of the CHM13 telomeres lack adjacent TAR1 repeats is unknown.

Overall, our results demonstrate the first single-molecule and single-nucleotide resolution maps of the genetic and chromatin component of all human telomeres, providing insights into the structure and function of human telomeres as well as their somatic stability.

## Acknowledgments

We thank Evan Eichler for providing CHM13 cells and for his helpful comments. We also thank Duncan Baird, Glennis Logsdon, Stephanie Bohaczuk, Jay Sarthy, and John Stamatoyannopoulos for their helpful comments and feedback. We are grateful to Katherine Munson for assistance with PacBio sequencing. This work was facilitated through the use of advanced computational, storage, and networking infrastructure provided by the Hyak supercomputer system and funded by the STF at the University of Washington. We thank the Telomere-to-Telomere (T2) consortium for generating a complete reference genome of CHM13 cells.

## Authors contributions

A.B.S., J.R. and B.M. designed and performed the experiments. A.B.S., D.D., A.S.C., and T.R. performed the computational analyses. A.B.S., D.D., A.S.C., J.R. and T.R. wrote the manuscript.

## Data and materials availability

The datasets generated during the current study will be available upon publication at the GEO repository (GEO accession GSE186009). The code used to generate the datasets used in this study will be available upon publication.

## Supplementary Materials

### Materials and Methods

#### Single-molecule chromatin fiber sequencing of CHM13 cells

CHM13 cells were grown in AmnioMax C-100 Basal Medium (Invitrogen 12558011), which includes the AmnioMAX™ C-100 Supplement to approximately 90% confluency. Growth media was changed daily and cells were split using 0.25% trypsin (Gibco 25200056). One million CHM13 cells per sample were pelleted at 250 x g for 5 minutes (4 samples were processed in parallel), washed once with PBS, and then pelleted again at 250 x g for 5 minutes. Each cell pellet was resuspended in 60 μL Buffer A (15 mM Tris, pH 8.0; 15 mM NaCl; 60 mM KCl; 1mM EDTA, pH 8.0; 0.5 mM EGTA, pH 8.0; 0.5 mM Spermidine) and 60 μL of cold 2X Lysis buffer (0.1% IGEPAL CA-630 in Buffer A) was added and mixed by gentle flicking then kept on ice for 10 minutes. Samples were then pelleted at 4°C for 5 min at 350 x g and the supernatant was removed. The nuclei pellets were gently resuspended individually with wide bore pipette tips in 57.5 μL Buffer A and moved to a 25°C thermocycler. 1 μL Hia5 MTase (200 U)^18^ and 1.5 μL 32 mM S-adenosylmethionine (NEB B9003S) (0.8 mM final concentration) were added, then carefully mixed by pipetting the volume up and down 10 times with wide bore tips. The reactions were incubated for 10 minutes at 25°C then stopped with 3 μL of 20% SDS (1% final concentration) and transferred to new 1.5 mL microfuge tubes. The sample volumes were adjusted to 100 μL by the addition of 37 μL PBS. To that, 500 μL of HMW Lysis Buffer A (Promega Wizard HMW DNA Extraction Kit A2920) and 3 μL RNAse A (ThermoFisher EN0531) were added. The tubes were mixed by inverting 7 times and incubated 15 minutes at 37°C. 20 μL of Proteinase K (Promega Wizard HMW DNA Extraction Kit A2920) was added and the samples mixed by inverting 10 times and incubated 15 minutes at 56°C followed by chilling on ice for 1 minute. Protein was precipitated by the addition of 200 μL Protein Precipitation Solution (Promega Wizard HMW DNA Extraction Kit A2920). Using a wide bore tip, the samples were mixed by drawing up contents from the bottom of the tube and then expelled on the side of the tube 5 times. Tubes were centrifuged at 16,000g for 5 minutes. The supernatant was poured into a new 1.5 mL tube containing 600 μL isopropanol. This was mixed by gentle inversion 10 times, incubated for 1 minute at room temperature, then inverted an additional 10 times. DNA was precipitated by centrifugation at 16,000g for 5 minutes. The supernatant was decanted, the pellet washed with the addition of 70% ethanol, and the sample was centrifuged again at 16,000 g 5 minutes. After the supernatant was decanted, a quick spin was performed to facilitate removal of any residual ethanol. Open tubes were air dried on the bench for 15 minutes. The DNA was resuspended by the addition of 25 μL 10mM Tris pH 8.0. Tubes were stored overnight in 4°C and the next day were mixed gently by using a wide bore pipette tip, drawing up the sample 5 times and gently expelling the contents on the side of the tube. Samples were then stored at -80°C prior to library construction.

DNA shearing, library construction, and PacBio Sequel II sequencing were performed as previously described^18^ with the exception that 15-20kb fragments were targeted when shearing using the Megaruptor (Diagenode Diagnostics). In addition, we performed a high-pass size selection of the SMRTbell library using the Sage Science PippinHT platform (Sage Science cat. no. ELF0001) according to the manufacturer’s protocol, using a high-pass cutoff of 10-15 kb to target an average library size of 17-20 kb.

#### Single-molecule chromatin fiber sequencing of GM12878 cells

GM12878 cells were purchased from the Coriell Institute for Medical Research and were grown in RPMI 1640 with 2mM L-glutamine and 15% fetal bovine serum. Growth media was changed every 2-4 days and cells were maintained at a density between 2-10x10^5^ cells/mL. GM12878 nuclei were isolated similarly to above with the exception that the 2X Lysis buffer contained 0.05% IGEPAL CA-630 in Buffer A. Fiber-seq reaction conditions, DNA isolation, shearing, and PacBio library construction were performed as described above for CHM13 cells.

#### Single-molecule chromatin fiber sequencing of K562 cells

K562 cells were grown and nuclei isolated as previously described^18^. Fiber-seq reactions were performed using the following conditions: (1) 200 Units of Hia5 for 10 minutes at 25°C; (2) 500 Units of Hia5 for 10 minutes at 25°C; (3) 1500 Units of Hia5 for 10 minutes at 25°C; (4) 200 Units of Hia5 for 10 minutes at 16°C; (5) 500 Units of Hia5 for 10 minutes at 16°C; or (6) 1500 Units of Hia5 for 10 minutes at 16°C. The same reaction buffer and nuclei amount was used as described above. The reactions were all stopped as above and DNA isolation, shearing, and PacBio library construction were performed as described above for CHM13 cells.

#### Hia5 expression and *in vitro* MTase activity assessment

Hia5 m6A-MTase was purified as previously described^18^, resulting in an enzyme concentration of 200 Units per μL, with a single unit of Hia5 defined as the maximum dilution of Hia5 that still results in near complete digestion by DpnI of Hia5-treated, purified dsDNA PCR product that contains a GATC motif. *In vitro* Hia5 m6A-MTase activity was performed as previously described^18^ with the following modifications: 60 μL reactions contained 1 μg of the dsPCR product substrate DNA, and various concentrations of Hia5 m6A-MTase [200 Units, 500 Units, or 1,500 Units] in Buffer A (15 mM Tris, pH 8.0; 15 mM NaCl; 60 mM KCl; 1mM EDTA, pH 8.0; 0.5 mM EGTA, pH 8.0; 0.5 mM Spermidine) supplemented with 0.8 mM S-adenosyl- methionine (NEB B9003S). A negative control was prepared without MTase. The reactions were mixed by gentle flicking of the PCR strip tubes and a quick spin down before a 1 hour incubation at either 25°C, or 16°C. DpnI digestion and electrophoresis conditions were as previously described^18^.

#### Mapping single-molecule chromatin fibers to CHM13 T2T genome

3 Sequel II SMRT cells for CHM13 Fiber-seq, 2 Sequel II SMRT cells for GM12878 Fiber-seq, and 1 multiplexed Sequel II SMRT cell for K562 Fiber-seq were processed individually. Using the raw subread bam files and the CHM13 v1.0 reference genome, we ran the findMethylationPipeline_no_probability.sh script on a SLURM managed cluster (***see data and code availability below***). This pipeline takes raw PacBio subread bam files and converts them into a comma separated format with per-base IPD ratio information for each read. Specifically, as part of this script, CCS reads were first generated using *pbccs (v6.0.0) (*https://ccs.how/) with the --chunk flag to parallelize across multiple nodes. Next, CCS reads were aligned to the CHM13 genome build v1.0 using *pbmm2 (v 1.2.0)* with the -N 1, --preset CCS, and --sort flags. This script then extracted the specific molecular identifier (ZMWID) for each aligned CCS read and the genomic sequence to which the CCS read aligned. It then iteratively used *bamsieve (v 0.2.0)* to extract all subreads with the same ZMWID using the --whitelist flag. Extracted subreads were then aligned to the genomic coordinates to which that ZMWID CCS moleculevmapped using *pbmm2 (v 1.2.0)* and flags: --sort, --preset SUBREAD. We then run *ipdSummary (v 3.0)* (https://github.com/PacificBiosciences/kineticsTools/blob/master/kineticsTools/ipdSummary.py) with the CCS-specific extracted genomic sequence as the --reference with these flags: --pvalue --identify m6A --csv <csv_file_name.csv> --numWorkers <number of available threads>. Finally, SMRT cell IDs were appended to each ZMWID to ensure that each molecule had a unique identifier when combining data across multiple SMRT cells.

Once findMethylationPipeline_no_probability.sh has completed, all ZMWID *ipdSummary* CSVs from a single SMRT cell were concatenated into a single file.

Next, using the *ipdSummary* CSV files from each SMRT cell, we trained a Gaussian Mixture Model (GMM) from adenine ipdRatios gathered from a subsample of 5,000 fibers by running *python3 faster_gmm.py --slurm input.csv*, similar to that previously described^20^. Once the GMM was trained on these 5,000 fibers, we applied the model to all fibers from a SMRT cell to calculate the methylation probability for each adenine contained within each fiber. Adenines with a m6A probability >= .99999999 were defined as being methylated.

To convert these methylation probabilities into bed files compatible with the UCSC browser, we ran *python3 new_csv_to_bed_with_model_apply.py -model trained_gmm.pkl input.csv output.bed*. This produces a bed12 formatted file that can be directly uploaded into the UCSC browser. Then we filtered out fibers devoid of m6A at the terminal ends by running *python3 cleanLowMethylation.py output.bed A_counts.txt* (To generate A_counts.txt we ran *awk -F”,” ‘$4==”A” { print $11}’ input.csv | sort | uniq -c > A_counts.txt*). Note that by convention, for each read in this file the first and final position of the read are given a size of ‘1’ so that it appropriately displays the ends of the read, independently of the methylation status at that position.

#### Identifying single-molecule nucleosome footprints

To identify nucleosomes along a single fiber, we used the *ipdSummary* CSV files and methylation bed12 files to train and apply a Hidden Markov Model (HMM)^44^ with a standardized post-hoc correction. Specifically, we ran the script *python3 hmmNucFinder_improved.py -A <output of build_A_bed.v2.py> -m <output of new_csv_to_bed_with_model_apply.py> -o <name of output HMM file>*. This script first projected methylation data into A/T space, using the *ipdSummary* CSV files to identify adenine positions along each fiber using the python3 script *build_A_bed.v2.py*. The HMM was then trained using a subsample of 5,000 fibers per SMRT cell and subsequently applied across all fibers from that SMRT cell. In tandem with the HMM delineation of nucleosomes, we also applied a simple finding approach that looked for unmethylated stretches larger than 85 bp in length (irrespective of A/T content). We used these ‘simple’ calls to refine our HMM nucleosome calls by splitting large HMM nucleosomes that contained multiple ‘simple’ calls. We also refined the terminal boundaries of each nucleosome by bookending it to the nearest m6A call within 10bp, if present. This produces a bed12 formatted file that can be directly uploaded into the UCSC browser. Note that by convention, for each read in this file the first and final position of the read are given a size of ‘1’ so that it appropriately displays the ends of the read, independently of the nucleosome footprint status at that position.

#### Quantifying single-molecule m6A-MTase accessibility

To identify single-molecule patches of m6A-MTase accessibility (MTase-sensitive patches - MSPs), we took the inverse of the nucleosome positions across single fibers with the script invertBed.py (*python3 invertBed.py bed_file > inverted_bed_file).* This produces a bed12 formatted file that can be directly uploaded into the UCSC browser. Note that by convention, for each read in this file the first and final position of the read are given a size of ‘1’ so that it appropriately displays the ends of the read, independently of the MSP status at that position.

#### Computational filtering of telomeric Fiber-seq reads

To only include high quality reads in our analysis we developed specific filters for reads from CHM13 cells and GM12878 cells. To filter for high quality CHM13 reads we required that 80% of the terminal telomeric 500bp on each read have a Phred Quality score >= 30. For GM12878 reads, we required that 80% of the terminal telomeric 500bp on each read have a Phred Quality score >= 30, have a mapping quality score (MAPQ) >= 40, and was not a secondary alignment. For the GM12878 analysis we focused on fibers from these telomere ends, as the coverage at these telomeres was consistent with genome-wide coverage of GM12878, indicating that these telomere ends were appropriately assembled: 1p, 2p, 3p, 3q, 6q, 7q, 9q, 11p, 20p, 22p.

#### Identifying genetic architectures of each telomeric fiber

To identify the genetic architecture of telomeric fibers, we searched all 9-mers along each fiber’s sequence to label canonical repeats as 9-mers perfectly matching TTAGGGTTA, which is the minimal sequence for TRF1/2 occupancy. Following identification of all 9-mer locations, overlapping 9-mers were merged to generate runs of canonical vs. non-canonical sequence. We then merged this data with fiber-specific chromatin information to delineate the relationship between each fiber’s telomeric sequence and telomeric chromatin structure.

#### Generating consensus telomere genetic architectures

To generate per-telomere consensus genetic architectures we utilized the per-fiber genetic architectures we previously generated. To account for the fact that the genetic architectures were constructed from the read sequence and not genomic sequence, we used the base position that aligned to the telomere-subtelomere boundary as an anchor point for each fiber. With the molecular coordinates normalized between fibers, we then iterated through every position relative to the telomere/subtelomere boundary, aggregated the sequence classification (TTAGGG vs. Non-TTAGGG) of each fiber at that position, and then classified into these 4 categories :

1. Germline canonical: Position relative to subtelomere is also called as canonical in >=50% of fibers at that position.
2. Somatic canonical: Position relative to subtelomere is also called as canonical in <50% of fibers at that position.
3. Germline non-canonical: Position relative to subtelomere is also called as non-canonical in >=50% of fibers at that position.
4. Somatic non-canonical: Position relative to subtelomere is also called as non-canonical in <50% of fibers at that position.

#### Identifying the nucleosomal architecture of telomeric reads

To properly identify the nucleosomal architecture of telomeric reads we modified our existing Fiber-seq pipeline to account for mapping difficulties that may arise from somatic alterations in the telomere repeat DNA content. First, we aligned CCS reads to the CHM13 genome (v1.0) and extracted ZMWIDs for all reads aligning to the telomeric sequence. Next, we modified the Fiber-seq computational pipeline to align each fiber to itself rather than the CHM13 genome (referred to as a CCS-based m6A profile as opposed to a genome reference-based m6A profile), enabling us to identify m6A patterns that may be unique to the DNA content of each read. To set these self-aligned fibers in a CHM13 coordinate space, we used their CCS alignment to the CHM13 genome to identify the centromeric end of the telomere TTAGGG repeat. We then set this position as 0 in the CCS-based m6A profile, allowing us to relate each individual fiber back to the CHM13 genome and to each other.

#### Quantifying the m6A methylation frequency of TTAGGG repeats genome-wide

To determine the m6A methylation frequency of the TTAGGG sequence, we first identified instances of the 7-mers CCTAACC, CTAACCC, and GTTAGGG in various genomic contexts and then calculated the proportion of these instances that contained an m6A at the central adenine. Note that these are the 7-mers corresponding to the three A/T base pairs in the TTAGGG repeat. Interstitial TTAGGG repeats were identified using repeat masker v2.0 file, which was downloaded from the UCSC T2T hub (http://t2t.gi.ucsc.edu/chm13/hub/t2t-chm13-v1.0/rmskV2/rmskV2.bigBed), and were defined as repeats containing iterations of TTAGGG or CCCTAA. These calculations were performed across a single SMRT cell.

#### Delineating CTCF binding elements and CTCF and cohesin occupancy

CTCF binding elements were identified genome-wide using the MA0139.1.meme motif from Jaspar. Specifically, the entire CHM13 v1.0 genome was scanned with this motif using *fimo* (version 4.11.2) with the --no-qvalue --thresh 0.001 --max-stored-scores 1000000 flags.

Identified elements that overlapped were resolved by selectively keeping the element with the lower p-value. To determine CTCF occupancy on each chromatin fiber we set these thresholds: (1) the motif of interest must contain 0 m6As within the binding element; (2) the motif must fall within a MSP >= 150 bp; and (3) the motif must have at least 3 m6As in the +/- 15bp bookending the motif instance. We applied these thresholds across each instance of an overlap between a motif and a fiber.

CTCF ChIP-seq raw FASTQ files were retrieved from 10 different cell lines (AG09309,AoAF, BJ, Caco-2, GM12878, HEEpiC, HepG2, K562, SAEC, SK-N-SH-RA) (<u>GSE30263</u>)^45, 46^. Reads were aligned to uniquely mapping locations along the CHM13 v1.0 genome using bwa mem (v0.7.17-r1188). Aligned reads were sorted and indexed using samtools (v1.11). ChIP-seq occupancy peaks were identified for each replicate with their respective control, using macs2 callpeak with a q-value of 0.01 (v2.2.7.1). Peaks were sorted by p-value (-log10) and the replicates were then assessed for consistency and reproducibility by calculating the irreproducibility discovery rate (IDR), using idr with a threshold of 1% (v2.0.4.2). Signal tracks were generated by calculating the fold-enrichment and log likelihood from pileup bedGraph files with macs2 bdgcmp, removing off-chromosome lines using bedTools slop (v2.30.0) + UCSC bedClip (v302.1 kentUtils) and transforming the resulting clip files to bigWig format.

ChIP-seq raw FASTQ files were retrieved for treated and untreated samples: RAD21, SMC1, CTCF, NIPBL, and input (GSM2809609 to GSM2809614, GSM2809637, GSM2809638, GSM3242972 and GSM3242973), all retrieved from the GEO accession series GSE104888^47^. All runs from the same sample were merged prior to alignment. All samples (single and paired-end) were aligned to the CHM13 V1.0 reference genome with bwa-mem -t 40 (v.0.7.17-r1188), sorted and indexed using samtools (v1.11). Peaks were called using macs2 callpeak (v.2.2.7.1) with parameters: -f BAM, -B, –SPMR, -q 0.01. Peaks were sorted by -log10 (p-value). The output bedgraph files were used to retrieve signal data and narrowPeak files used to retrieve peak data.

#### CHM13 transcriptional activity

Iso-seq and RNA-seq data from CHM13 cells were downloaded from the T2T repository^17^. For RNA-seq, FASTQ files from both libraries were merged. CHM13 liftoff based GENCODE gene models (http://t2t.gi.ucsc.edu/chm13/hub/t2tChm13_20200727/liftOffGenes/) were transformed to GTF format using the UCSC bigGenePredToGenePred and genePredToGtf tools (v302.1 kentUtils). An index was generated for the CHM13 v1.0 genome using the STAR algorithm (v2.7.8a) with the following flags: --runMode genomeGenerate --sjdbOverhang 74. Next, reads were aligned with STAR using the following flags: --runMode alignReads, --twopassMode Basic, --alignSoftClipAtReferenceEnds Yes, --quantMode TranscriptomeSAM GeneCounts, -- quantTranscriptomeBAMcompression -1,--outSAMtype BAM SortedByCoordinate,-- outBAMcompression -1,--outSAMunmapped Within,--genomeLoad NoSharedMemory. We calculated the total number of RNA-seq reads in the terminal 250kb of each chromosome, with the first bin starting at the end of the chromosome.

CHM13 Iso-Seq data was downloaded from the NCBI sequence read archive (SRA) accession SRR12519035. Iso-Seq bam files were aligned to CHM13 v1.0 using *pbmm2 (v1.2.0) --preset ISOSEQ.* Aligned bam files were subsequently converted to bed format. To investigate differences between TAR1 and non-TAR1 containing terminal ends we intersected the respective non-telomeric terminal 500-kb with the IsoSeq bed file and then counted the number of intersections for each terminal end with the following command : *bedtools intersect -wa -a respective.terminal.region.bed -b isoseq.aligned.sorted.bed | sort | uniq -c.* Terminal ends with no transcripts aligning were given a value of 0.

#### TAR1 repeat clustering

TAR1 repeats were identified using repat masker v2.0 file, which was downloaded from the UCSC T2T hub (http://t2t.gi.ucsc.edu/chm13/hub/t2t-chm13-v1.0/rmskV2/rmskV2.bigBed), and were filtered for those with a size >=500bp. A FASTA file containing DNA oriented to the TAR1 repeat orientation was generated for each of these elements using bedtools getfasta and the CHM13 v1.0 reference genome. The FASTA sequences were then aligned using the multiple sequence aligner program, MAFFT^48^ using the following flags: --reorder --maxiterate 1000 -- thread 16 . The resulting multiple sequence alignment was used to create a maximum likelihood phylogenetic tree with the tool RAxML (Stamatakis, 2014), by running *raxmlHPC-PTHREADS - f a -p 12345 -x 12345 -s {TAR1 elements MSA fasta} -m GTRGAMMA -# 100 -T 8 -n TAR1*.

**Fig. S1.**
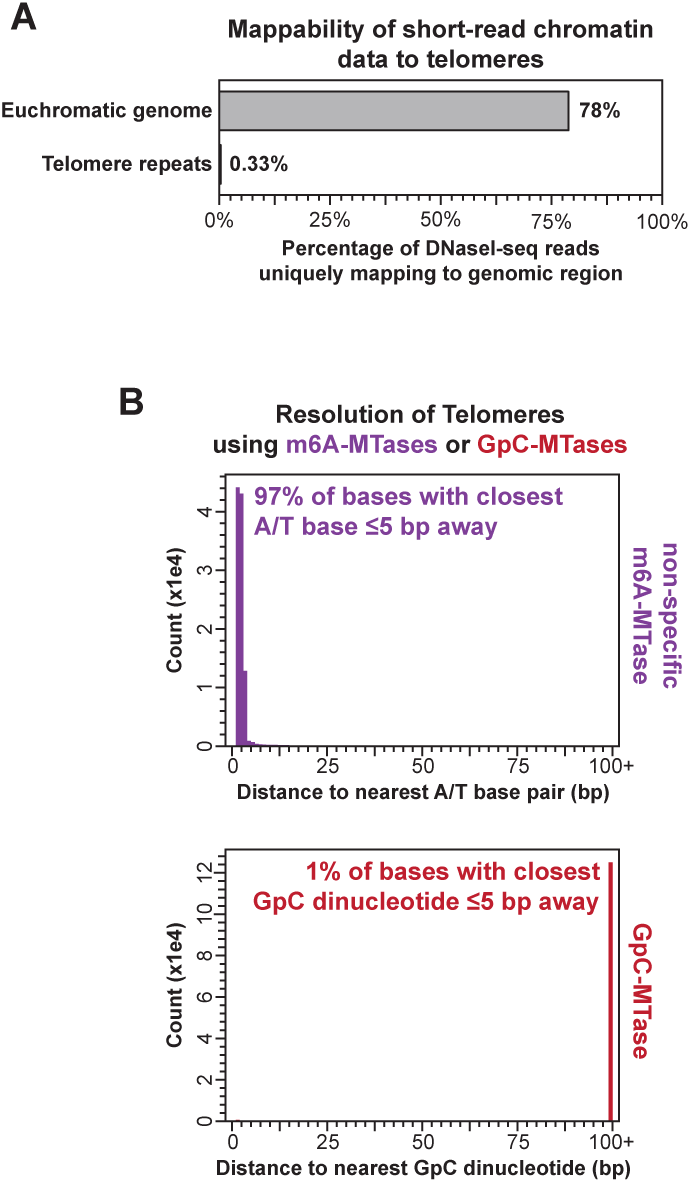
Resolution of short-read and long-read epigenomic mapping methods. **(A)** Percentage of reads from GM12878 DNaseI seq experiment uniquely mapping to various segments of the genome. Euchromatic genome defined as genomic regions not contained within repeats. **(B)** Distance between neighboring A/T base pairs or GC dinucleotides outside of a CpG context within telomeres.

**Fig. S2.**
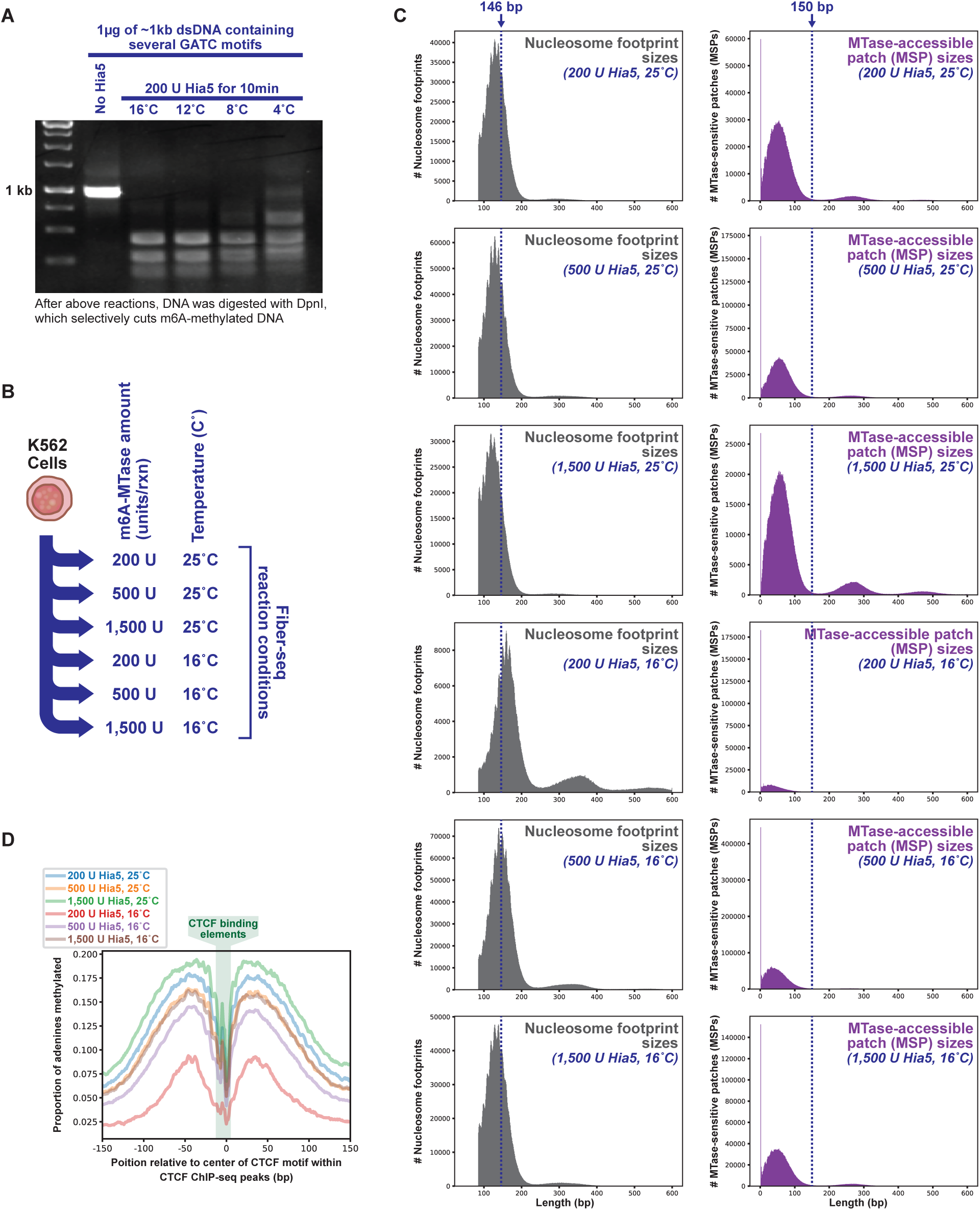
Optimization of Fiber-seq for human samples. **(A)** DNA gel of Hia5 m6A-MTase activity at various temperatures. Specifically, 1μg of ∼1kb dsDNA PCR product that contains several GATC motifs was treated with 200 units of Hia5 for 10 minutes at various temperatures, or untreated with Hia5. The DNA was then purified and digested with DpnI, which will selectively cut DNA molecules with methylated GATC motifs. **(B)** Schematic for testing optimal Fiber-seq reaction conditions in human cells. Specifically, all reaction were performed for 10 minutes with the same number of cells, and the temperature and enzyme concentration were varied. **(C)** Histograms of (left) nucleosome footprint sizes and (right) MTase-accessible patch (MSP) sizes from Fiber-seq reactions performed with various amounts of enzyme or at various temperatures. **(D)** Aggregate m6A profile surrounding ChIP-seq positive CTCF binding elements from Fiber-seq libraries obtained according to panel b.

**Fig. S3.**
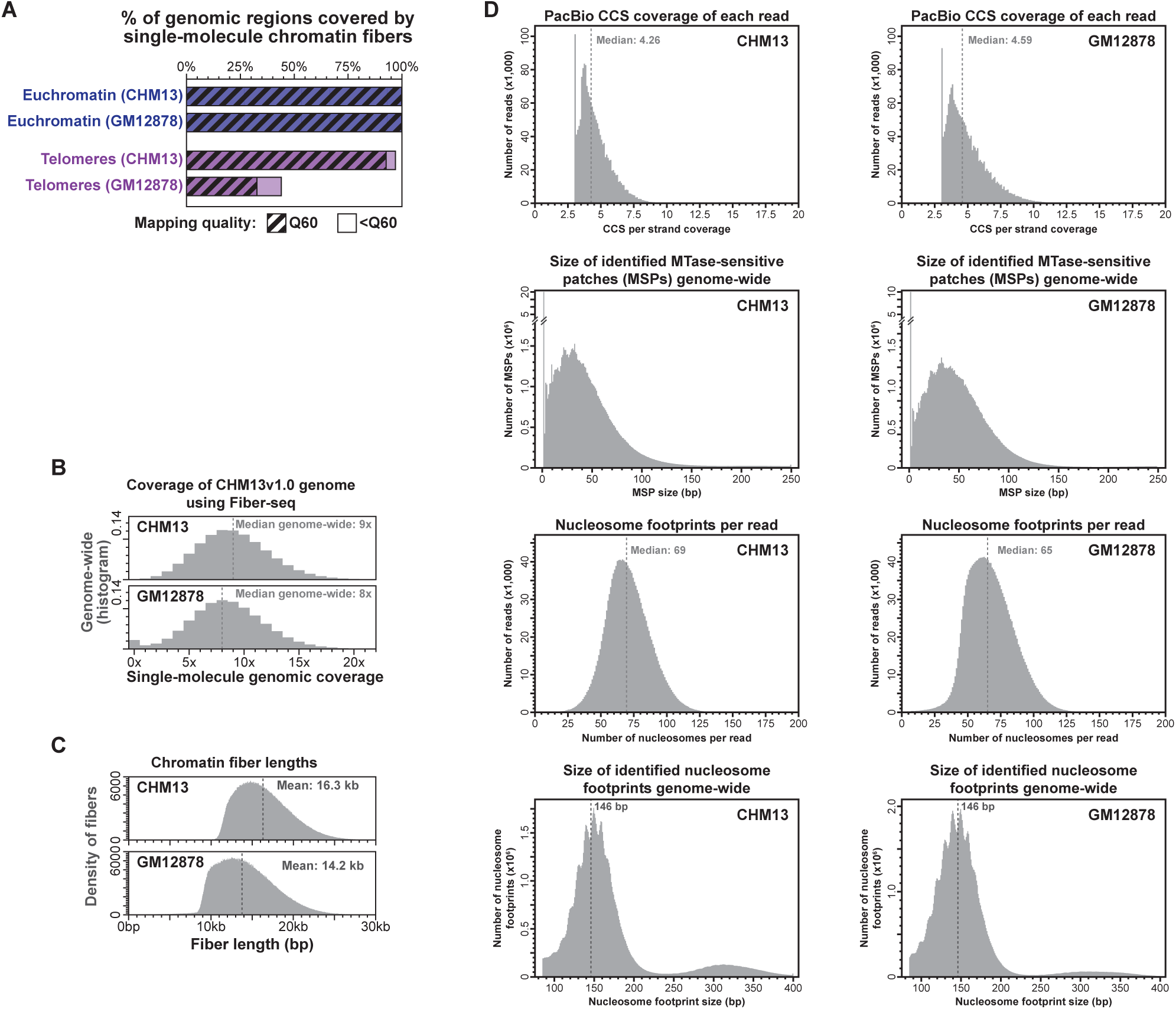
Fiber-seq statistics for CHM13 and GM12878 cells mapped to CHM13 genome. **(A)** Mapping quality and coverage of various regions of the CHM13 v1.0 genome obtained by performing Fiber-seq on GM12878 cells or CHM13 cells. **(B)** Coverage of the CHM13 v1.0 genome obtained by performing Fiber-seq on GM12878 cells or CHM13 cells. **(C)** Histogram showing the size distribution of chromatin fibers sequenced using Fiber-seq from GM12878 and CHM13 cells. **(D)** Histograms of PacBio circular consensus sequencing (CCS) coverage of each strand, the number of nucleosome footprints identified on each fiber, the size of MSPs identified, and the size of nucleosome footprints identified for reads from the GM12878 and CHM13 Fiber-seq experiment.

**Fig. S4.**
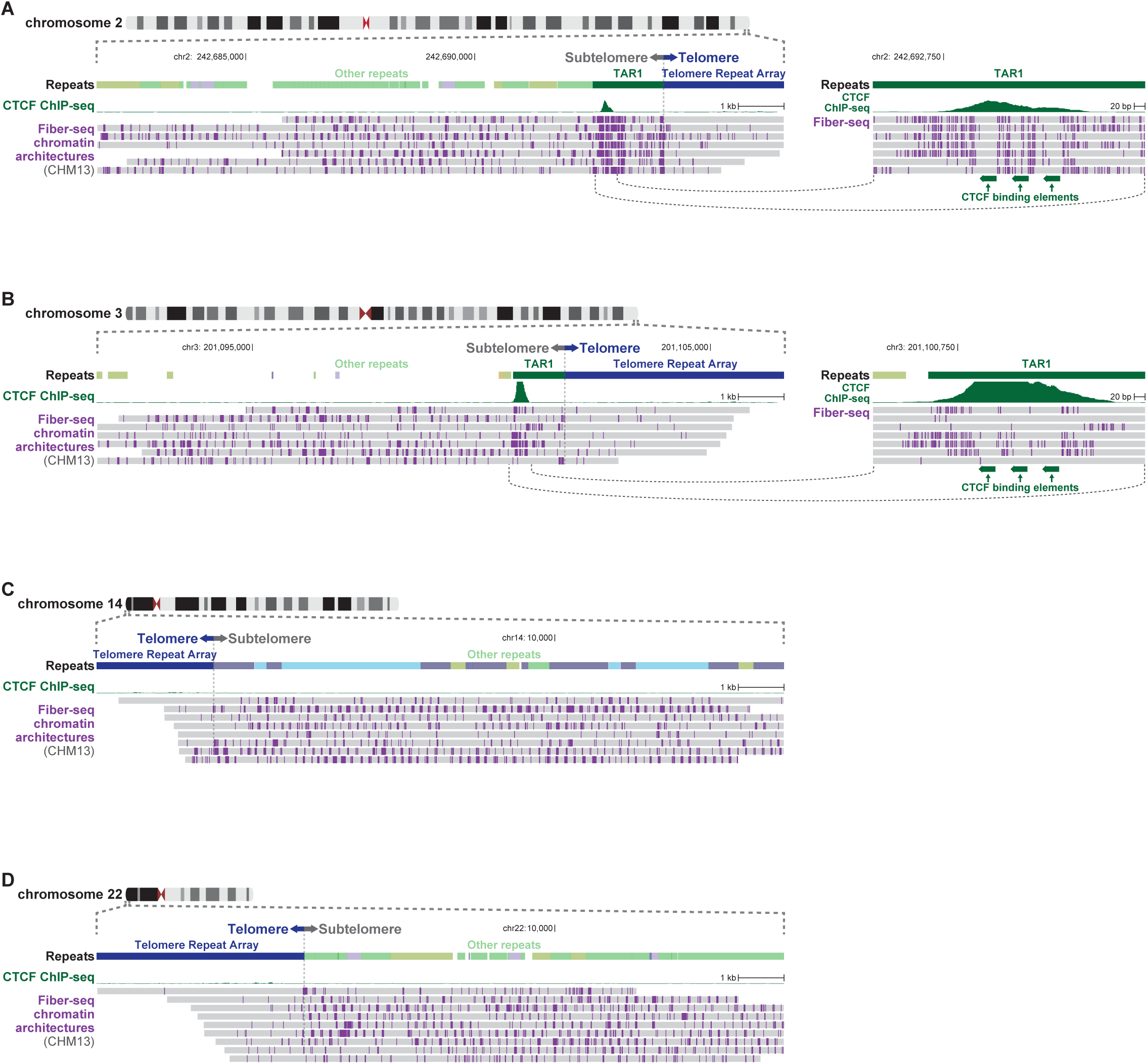
CHM13 telomere chromatin architecture (A-D) Genomic loci showing per-molecule genetic and chromatin architectures for all fibers from (A) 2q, (B) 3q, (C) 14p, and (D) 22p telomeres.

**Fig. S5.**
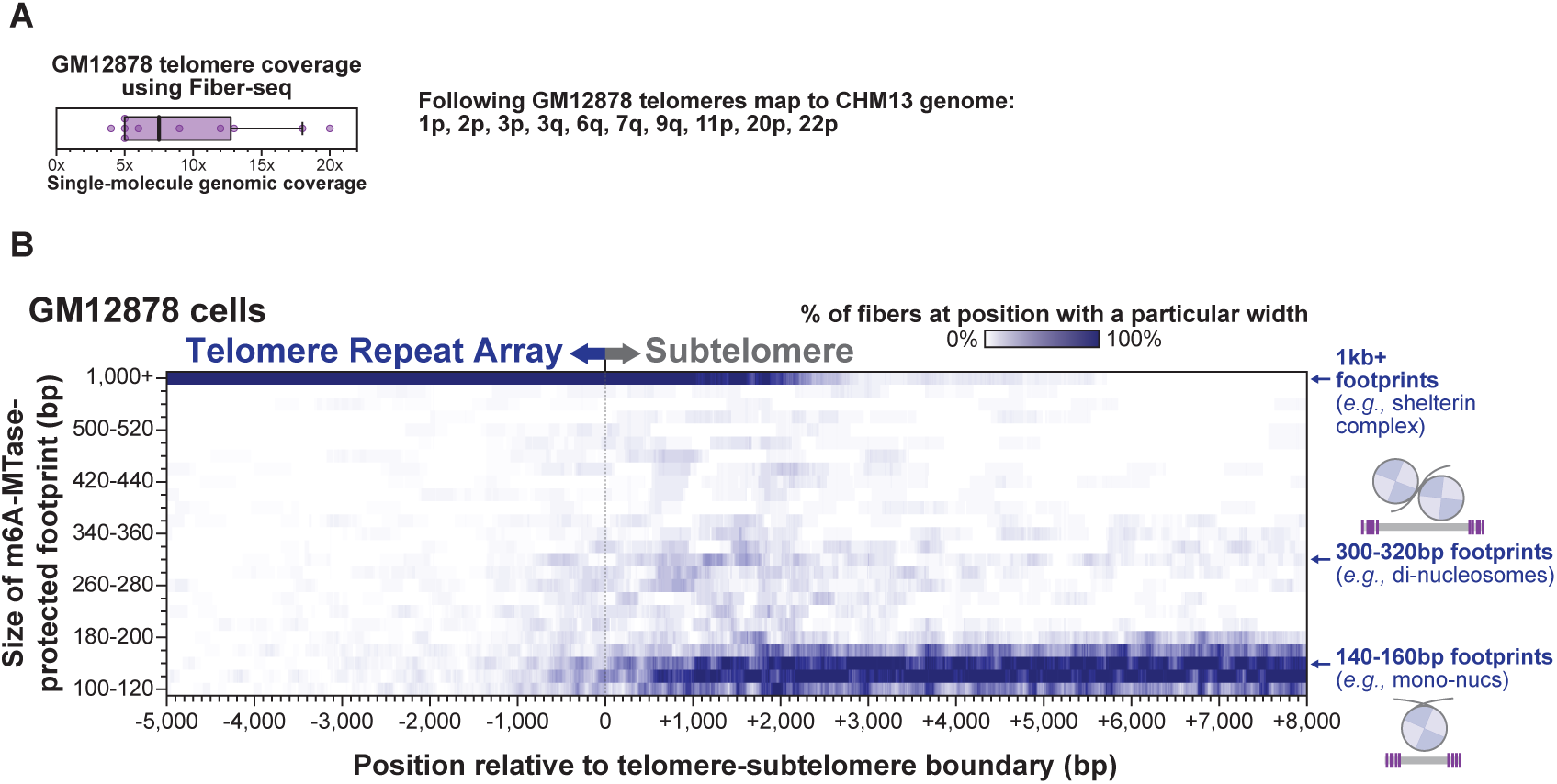
GM12878 telomere chromatin architecture. **(A)** Coverage of single-molecule chromatin fibers along individual GM12878 telomeres, as well as the list of telomeres that were appropriately mapped using GM12878 Fiber-seq and the CHM13 reference genome. **(B)** Heatmap showing the distribution of footprint sizes relative to the telomere-subtelomere boundary across these GM12878 telomeres.

**Fig. S6.**
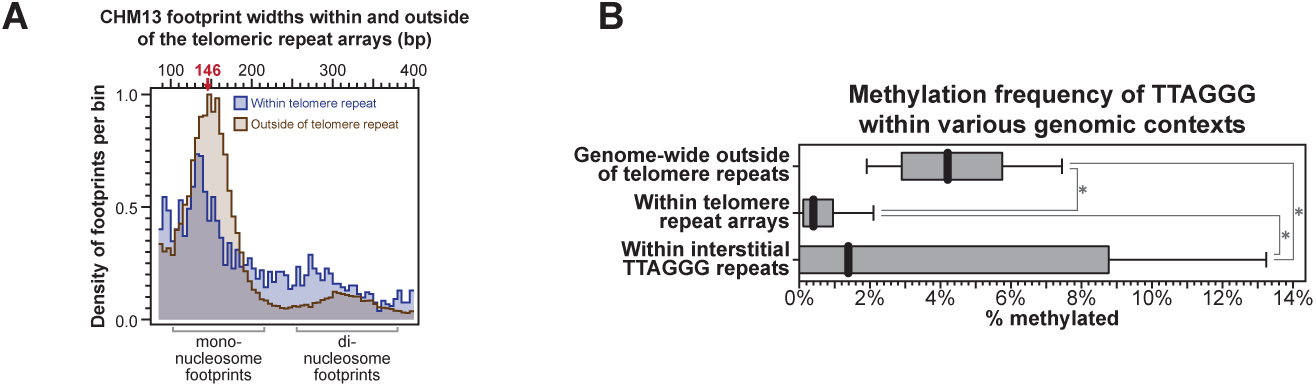
Nucleosome and methylation architecture of telomere and non-telomere DNA. **(A)** Histogram of nucleosome footprint widths within telomere repeat arrays (blue), and genome-wide (tan) from CHM13 cells. **(B)** Methylation frequency of the central adenine in CTAACCC 7-mers located within various genomic contexts.

**Fig. S7.**
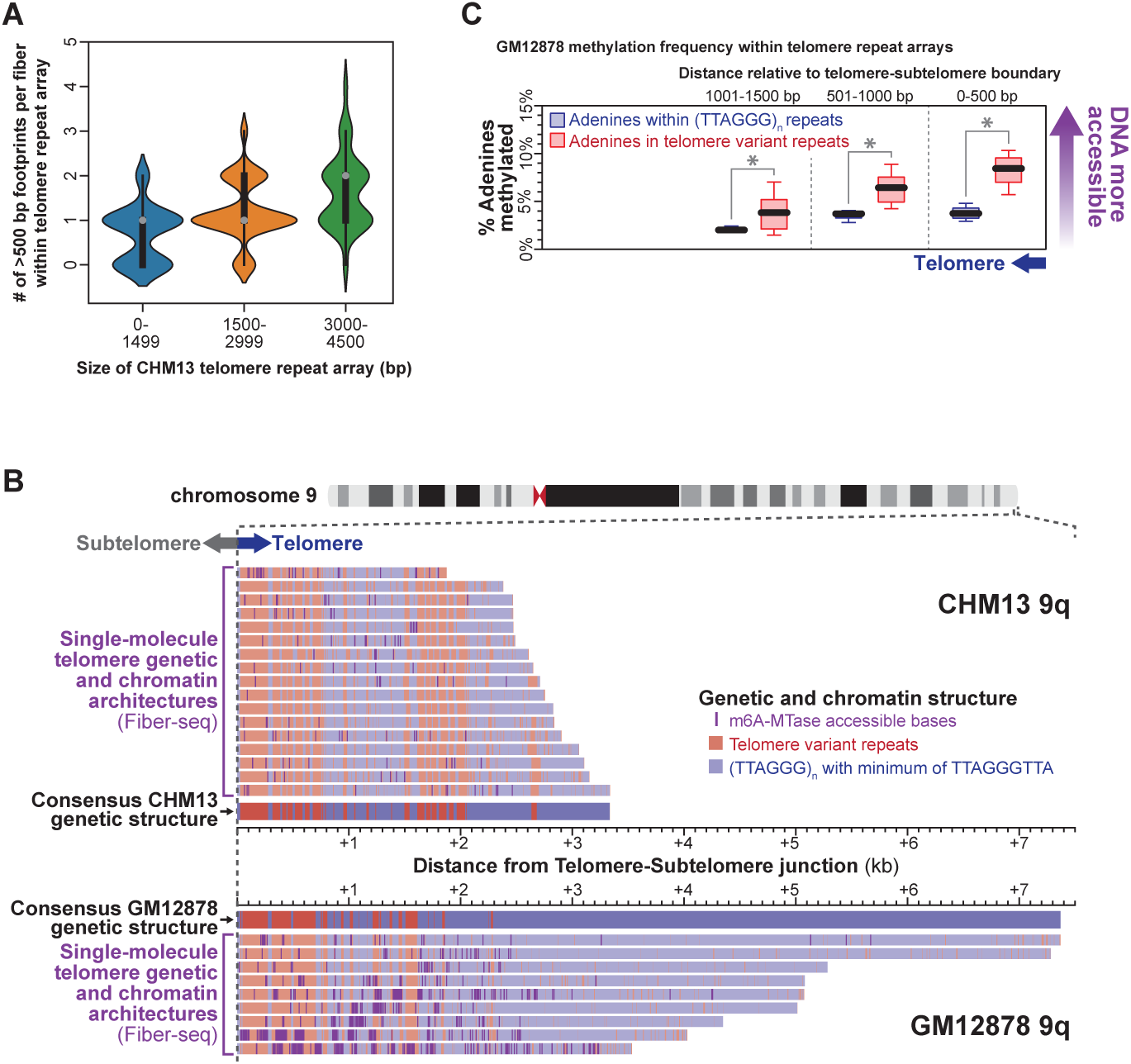
GM12878 and CHM13 telomere chromatin and genetic architectures. **(A)** Violin plots showing the number of >500 bp footprints per fiber within CHM13 telomere repeat arrays based on the size of the telomere repeat array. **(B)** Genomic locus showing per-molecule genetic and chromatin architectures for all fibers overlapping the chromosome 9q telomere in CHM13 and GM12878 cells. Location of per-fiber TTAGGG repeats, and telomere variant repeats in blue and red, respectively. Consensus sequence of CHM13 and GM12878 9q telomeres shown as well. **(C)** m6A-methylation frequency of TTAGGG repeats and telomere variant repeats within GM12878 telomeres, relative to telomere-subtelomere boundary (* p- value <0.01 Mann-Whitney).

**Fig. S8.**
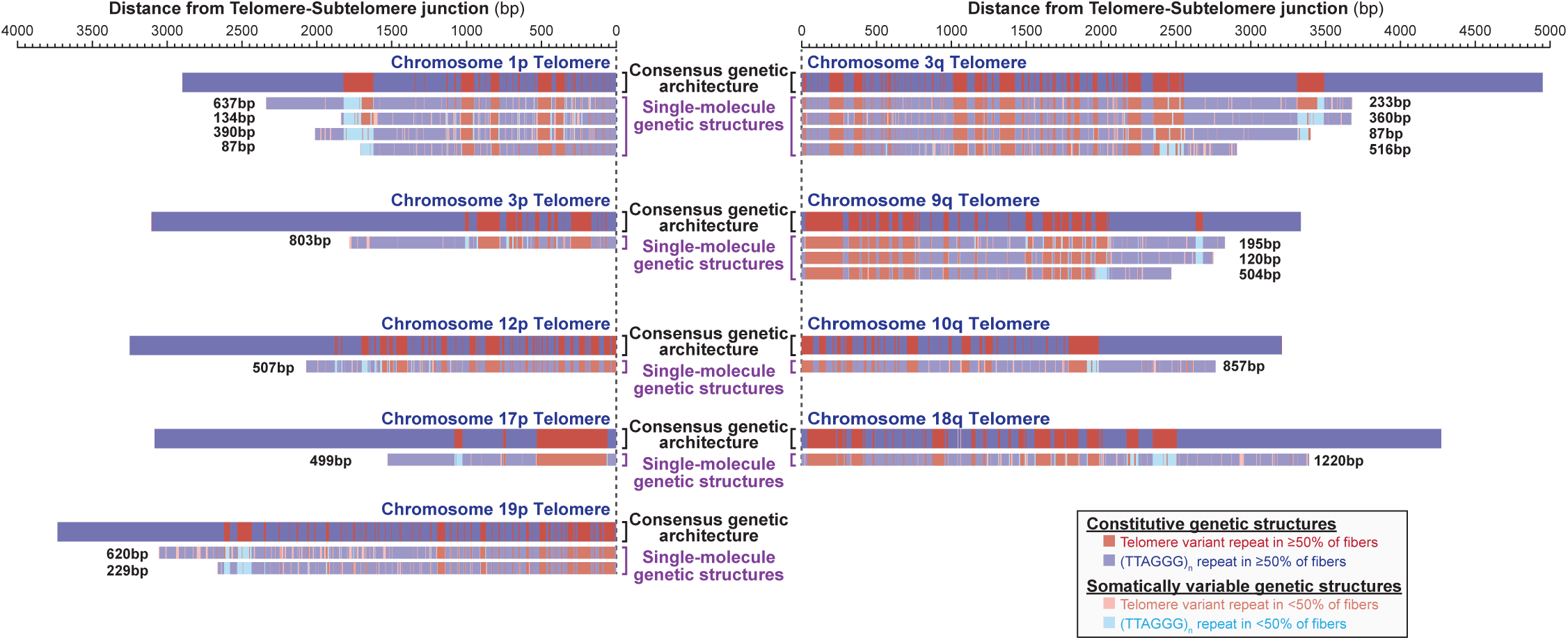
Somatic stability of CHM13 telomeres. Genomic loci showing per-molecule genetic architectures for all fibers that demonstrate somatically rearranged telomere variant repeats. Shown are the individual fibers that have undergone somatic reconstruction, as well as the consensus genetic architectures based on the other fibers from that telomere (these other fibers are not shown). Reads from telomeres 1p, 3p, 12p, 17p, 19p, 3q, 9q, 10q, and 18q are displayed, as these are the only reads that clearly demonstrated somatic reconstruction. Telomere fibers colored based on their sequence relative to the consensus sequence, with repeats matching the consensus in dark and repeats not matching the consensus in light colors. TTAGGG repeats, and telomere variant repeats on each fiber colored in blue and red respectively. The minimal length of each somatically restructured telomere end is displayed along side each fiber.

**Fig. S9.**
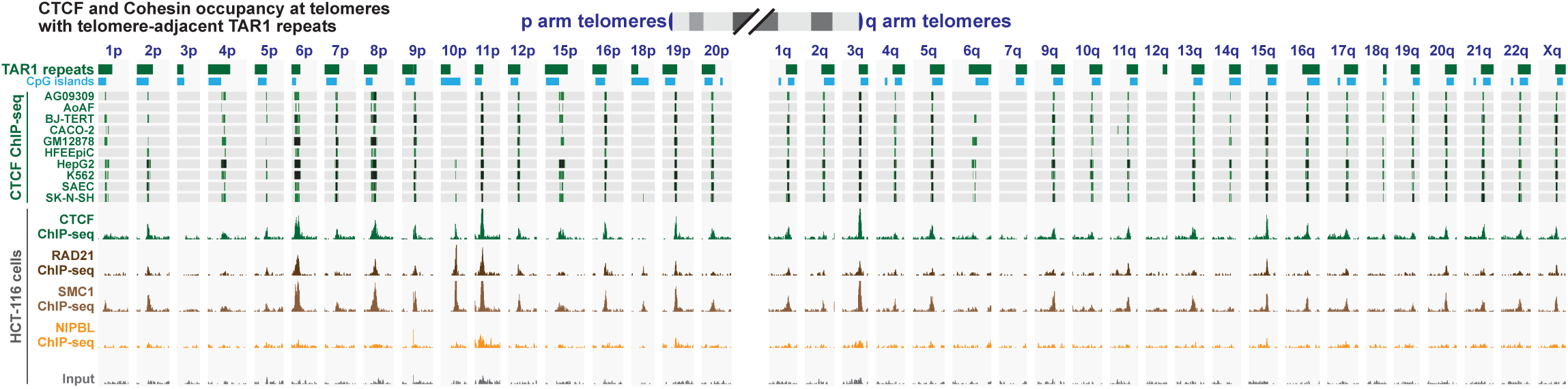
Chromatin architecture of TAR1 repeats. Genomic loci showing CTCF and Cohesin ChIP-seq signal from various cell lines mapped across the 39 telomere-adjacent TAR1 elements in the CHM13 genome. Track heights for all CTCF ChIP-seq samples, and HCT-116 cell samples are identical, respectively.

**Fig. S10.**
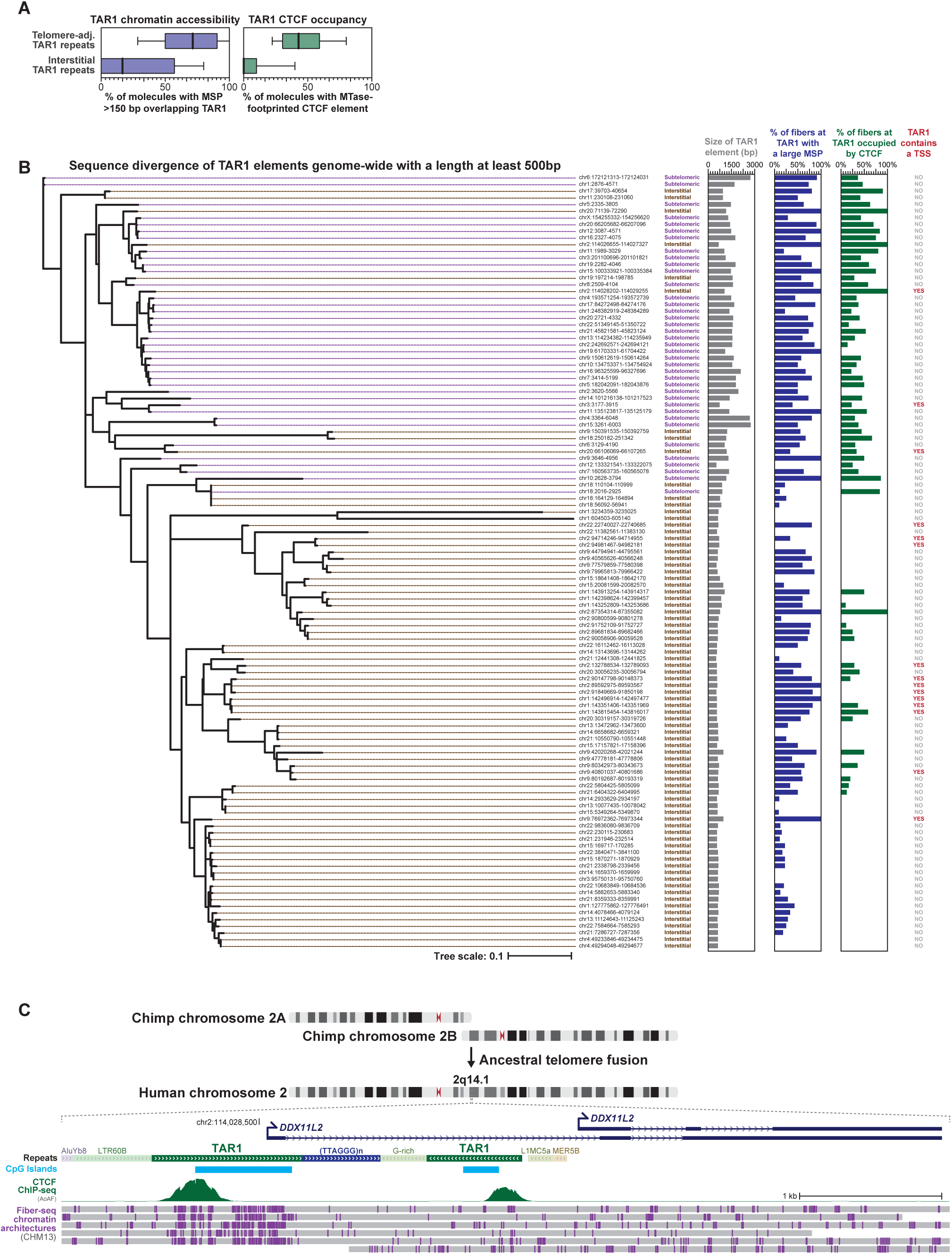
TAR1 repeat evolution and chromatin architecture. **(A)** Box-and-whisker plots for telomere-adjacent and interstitial TAR1 elements showing (left) the percentage of overlapping fibers at TAR1 repeat that contain an MSP>150bp, and (right) the percentage of overlapping fibers at TAR1 repeat that contain an occupied CTCF binding element. **(B)** Phylogenetic tree comparing the sequence of each of the interstitial and telomere-adjacent TAR1 repeats >500 bp in length. Displayed to the right are the sizes of each element, as well as the percentage of overlapping fibers containing an MSP>150 bp within the TAR1 repeat, or containing an occupied CTCF binding element within the TAR1 repeat. TAR1 repeat overlap with transcriptional start sites (TSS) is also displayed. **(C)** Genomic locus with an interstitial TAR1 element at the chromosome 2 telomere-telomere fusion site that functions as a promoter for a non-coding gene. Displayed is CTCF ChIP-seq signal, as well as single-molecule chromatin fiber architectures.

**Fig. S11.**
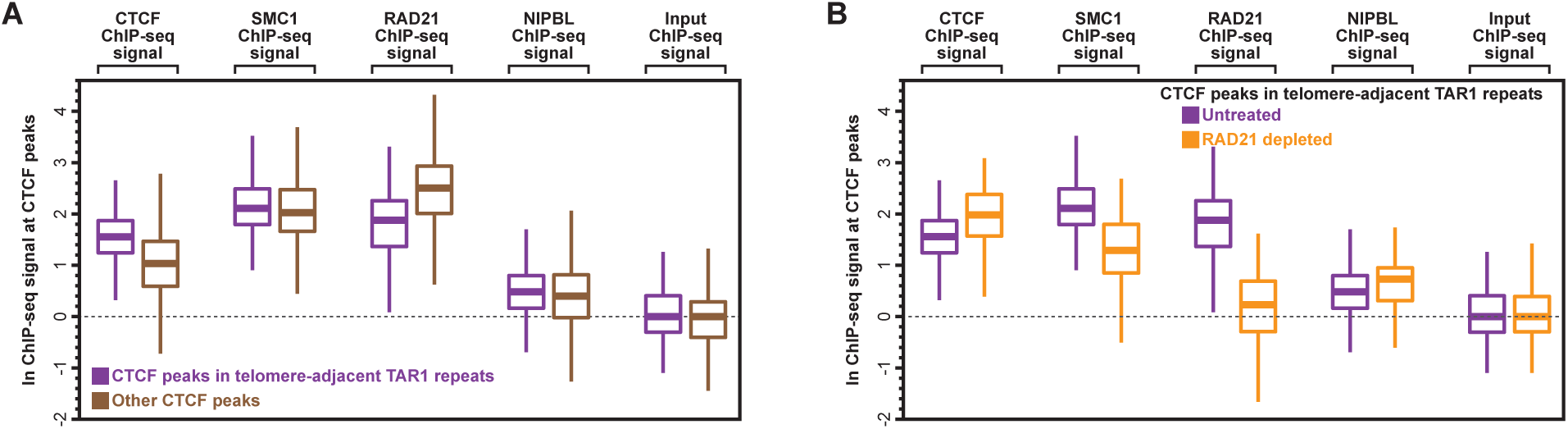
Cohesin signal at TAR1 repeat CTCF elements. **(A)** ChIP-seq signal for CTCF, cohesin components RAD21 and SMC1, cohesin loading factor NIPBL, and input at CTCF ChIP-seq peaks at telomere-proximal TAR1 elements, or other CTCF ChIP-seq peaks in HCT-116 cells. **(B)** ChIP-seq signal for CTCF, cohesin components RAD21 and SMC1, cohesin loading factor NIPBL, and input at CTCF ChIP-seq peaks at telomere-proximal TAR1 elements in HCT-116 cells before or after RAD21 depletion. ChIP-seq data obtained from Rao et al., *Cell* 2017, wherein RAD21 depletion was accomplished via an auxin-inducible degron (AID).

**Fig. S12.**
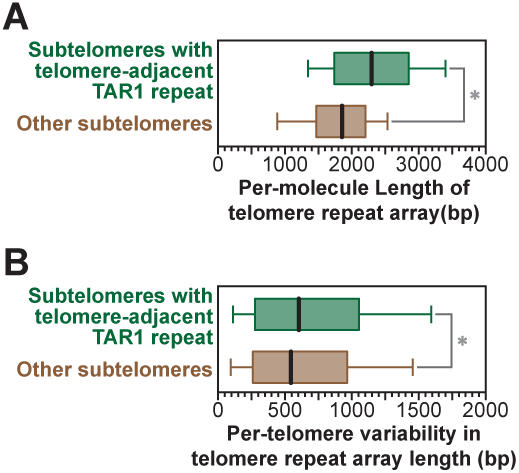
Telomere lengths at telomeres with adjacent TAR1 repeats in CHM13 cells. **(A)** Box-and-whisker plot showing length of telomere repeats for fibers from telomeres with telomere-adjacent TAR1 repeats. **(B)** Box-and-whisker plot showing the difference in fiber lengths from telomere fibers that originate from the same telomere (* p-value <0.01 Mann-Whitney).

